# Identification of PDE6D as a potential target of sorafenib via PROTAC technology

**DOI:** 10.1101/2020.05.06.079947

**Authors:** Yang Li, Qingshi Meng, Ping Wang, Xiaolei Liu, Qiuyu Fu, Yuting Xie, Yu Zhou, Xiangbing Qi, Niu Huang

**Affiliations:** National Institute of Biological Sciences, Beijing, No. 7 Science Park Road, Zhongguancun Life Science Park, Beijing, 102206, China; State Key Laboratory of Animal Nutrition, Institute of Animal Science, Chinese Academy of Agricultural Sciences, Beijing 100193, China; Tsinghua Institute of Multidisciplinary Biomedical Research, Tsinghua University, Beijing 102206, China

**Keywords:** PDE6D, PROTAC, Protein degradation, Ubiquitylation, Target identification

## Abstract

The identification of unknown target of a multi-kinase inhibitor sorafenib is important to better understand the mechanism of action of this drug in anti-cancer and anti-fibrotic treatments. Here, we report the combination of PROTAC technique with quantitative proteomic analysis to identify the unknown cellular targets of sorafenib. Sorafenib-based PROTAC can strongly degrade a non-kinase target PDEδ in different types of cells. We also confirmed the direct binding interaction of PDEδ with sorafenib by CETSA and SPR assays. Together, our research suggests that PDEδ is a new potential target of sorafenib, and PROTAC technology may be a promising approach for cellular target identification of bioactive compounds of interest.

## 1. Introduction

Sorafenib is a multi-kinase inhibitor and has been widely used in oncotherapy^1^. Sorafenib was reported against numerous protein kinases, including Raf-1 (IC_50_ = 6 nM), BRAF (IC_50_ = 22 nM), c-KIT (IC_50_ = 58 nM), FLT3 (IC_50_ = 58 nM), VEGFR2 (IC_50_ = 90 nM) and PDGFRβ (IC_50_ = 57 nM)^2^. In our previous study, we discovered sorafenib as a nanomolar ligand of non-kinase target 5-HT receptors (K_i_ =56 and 417 nM against 5-HT_2B_ and 5-HT_2C_, respectively) by virtual screening^3^. Recently, 5-HT_2B_ receptor was also considered as a promising target for the treatment of liver fibrosis^4^. Given sorafenib’s promiscuity, it is desirable to identify new unknown targets of sorafenib to better understand its molecular mechanism of action in anti-cancer and anti-fibrotic treatments.

Proteolysis Targeting Chimera (PROTAC) is a promising technology for degradation of target proteins by endogenous proteasomes, which comprises a ligand for the target of interest, conjugated to a ligand for an E3 ubiquitin ligase and a linker conjugating them^5^. By recruiting E3 ubiquitin ligase to target proteins through target-specific ligands, a broad range of target proteins that are dysregulated in cells have been degraded^6-15^. Typically, successful PROTACs were designed to degrade the specific cellular protein by using a high affinity ligand. However, multiple targets could be degraded simultaneously by using low affinity and promiscuous ligand-based PROTAC, such as P38α which could be degraded by the low binary binding affinity fore-tinib-based PROTAC^16, 17^. So far the formation and stabilization of the target:PROTAC:E3 ligase ternary complex were thought to be more important in determining the efficient degradation of the target^16, 18^. Since PROTAC does not require very high binding affinity to degrade potential targets, we are interested in applying the PROTAC technique to identify potential unknown targets of sorafenib.

Here, we report the design and synthesis of a sorafenib-based PROTAC T-S. Upon the recruitment of Cereblon (CRBN)/Cullin 4A E3 ligase complex, we identified the prenyl-binding protein PDEδ as a dominant target of sorafenib-based PROTAC T-S. Furthermore, we verified the direct binding interaction between sorafenib and PDEδ. These data gave us a new insight into the mechanism of action of sorafenib.

## 2. Results and discussion

### 2.1. Design of sorafenib-based PROTAC (PROTAC T-S)

We began our investigation by designing and synthesizing a sorafenib-based PROTAC. General synthetic routes to compound PROTAC T-S are provided in Scheme 1. The multi-kinase inhibitor sorafenib binds to its target kinases by forming two hydrogen bonds between its N-methylpicolinamide group and the ATP binding-site hinge region backbone^19^. To maximize the chance of finding the non-kinase targets of sorafenib, we coupled the linker with the N-methylpicolinamide group of sorafenib, and we hope that the linker added to the N-methylpicolinamide group would reduce the kinase binding activity. Since 3-polyethylene glycol (PEG) linkers were widely used in the literature for CRBN-recruiting PROTACs^15, 20, 21^, we coupled sorafenib with a 3-PEG linker conjugated to thalidomide (referred to PROTAC T-S) (Figure 1A). We then tested the kinase binding activity of PROTAC T-S by using known kinase targets of sorafenib, including BRAF, c-KIT, FLT3, P38α, VEGFR2 and PDGFRβ. PROTAC T-S at concentration 0.1 μM showed more than a two-fold decrease in inhibitory activity of those targets compared with sorafenib, except P38α (IC_50_ = 1.72 and 1.84 μM to sorafenib and PROTAC T-S, respectively) (Table S1).

**Figure 1.**
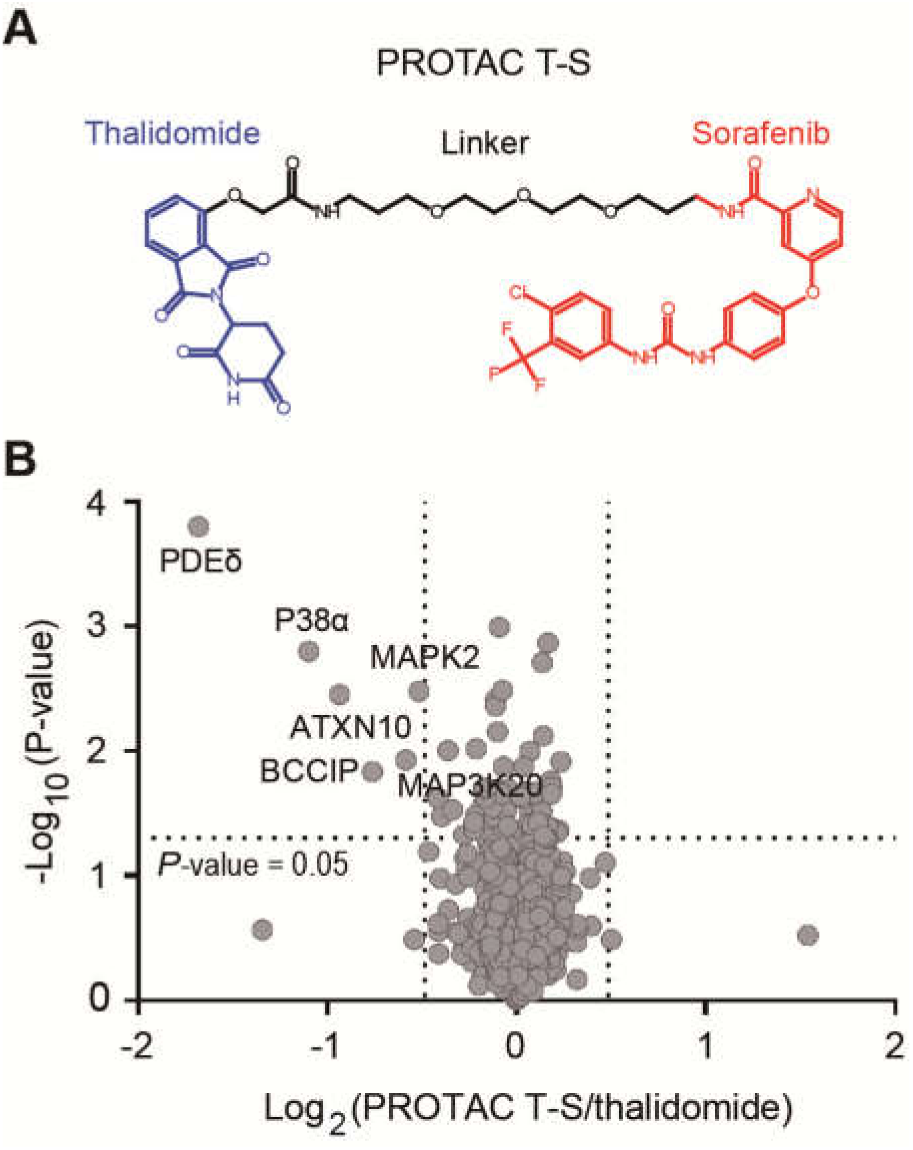
(A) Structure of sorafenib-based PROTAC. (B) A Tandem Mass Tags (TMT)-based mass spectrometry (MS) approach was used to assess the proteome changes in AML12 cells after 12 hours treatment of 2 μM PROTAC T-S or thalidomide. The identified total 5749 unique proteins were plotted as Log_2_ of fold change (PROTAC T-S/thalidomide) versus -Log_10_ of *p*-value (*t*-test).

### 2.2. PDEδ is a dominant target of PROTAC T-S

Moreover, among all chosen kinase targets (P38α, BRAF, RAF1 and c-Kit), P38α was the only one degraded by PROTAC T-S (Figure S1A). To further explore the potential targets degraded by PROTAC T-S, we applied a Tandem Mass Tag (TMT)-based quantitative proteomics approach to monitor the protein fold changes at the cellular level. Thalidomide was used as a negative control to exclude the undesired neo-substrates that recruited by thalidomide-bound CRBN^22^. We compared the proteomics results of AML12 cells after 12 hours of treatment with 2 μM PROTAC T-S or thalidomide. A total of 5749 unique proteins was identified in our experiments, but only 6 proteins were significantly down-regulated by more than 30% (*p*-value of 0.05) (Figure 1B). Among these six targets (PDEδ, P38α, ATXN10, BCCIP, MAP3K20, and MAPK2), three of them were non-kinases, which is according with our original designing of PROTAC T-S. It is notable that PDEδ was most efficiently degraded, with more than 80% decrease in protein expression after PROTAC T-S treatment compared to the thalidomide-treated group.

PDEδ was well known as a chaperone of some prenylated proteins^23^. The hydrophobic binding pocket of PDEδ binds the farnesyl group of KRAS4B and facilitates its membrane localization^24^. Blocking the PDEδ-KRAS interaction was considered as a promising target in cancer research, and novel PDEδ ligands bound to the prenyl-binding pocket were identified to inhibit RAS activation^25, 26^. Recently, a PROTAC version of a high-affinity PDEδ ligand deltarasin was developed to degrade PDEδ^27^. To confirm that the degradation of PDEδ detected by mass spectrometry (MS), we used Western blotting to analyze PDEδ protein levels in PROTAC T-S-treated AML12 cells. The degradation of PDEδ, unlike that of the known high affinity sorafenib-binding target Raf-1, in AML12 cells happens in a dose-dependent manner (Figure 2A). Sorafenib was reported to induce downregulation of some of its substrates via competition for chaperone Hsp90-Cdc37 binding^28^. To test whether Hsp90 chaperone system affect the stability of PDEδ, we used sorafenib alone to treat AML12 cells and found that only high dose of sorafenib can slightly reduce PDEδ levels (Figure S2). The degradation of PDEδ induced by PROTAC T-S was faster than the degradation of P38α, with more than half of PDEδ being degraded within one hour (Figure 2B and Figure S1B). To further assess the generality of PDEδ degradation mediated by PROTAC T-S, we used PROTAC T-S to treat different cell lines. PDEδ was efficiently degraded in all selected mice or human cell lines (Figure 2C and Figure S3). Furthermore, we used the quantitative real-time PCR to test whether PROTAC T-S downregulate PDEδ at the transcriptional level, the results showed that neither PROTAC T-S nor sorafenib-treated cells showed significant changes in PDEδ mRNA (Figure 2D). Thus, we determined that PDEδ is a novel target of sorafenib-based PROTAC T-S.

**Figure 2.**
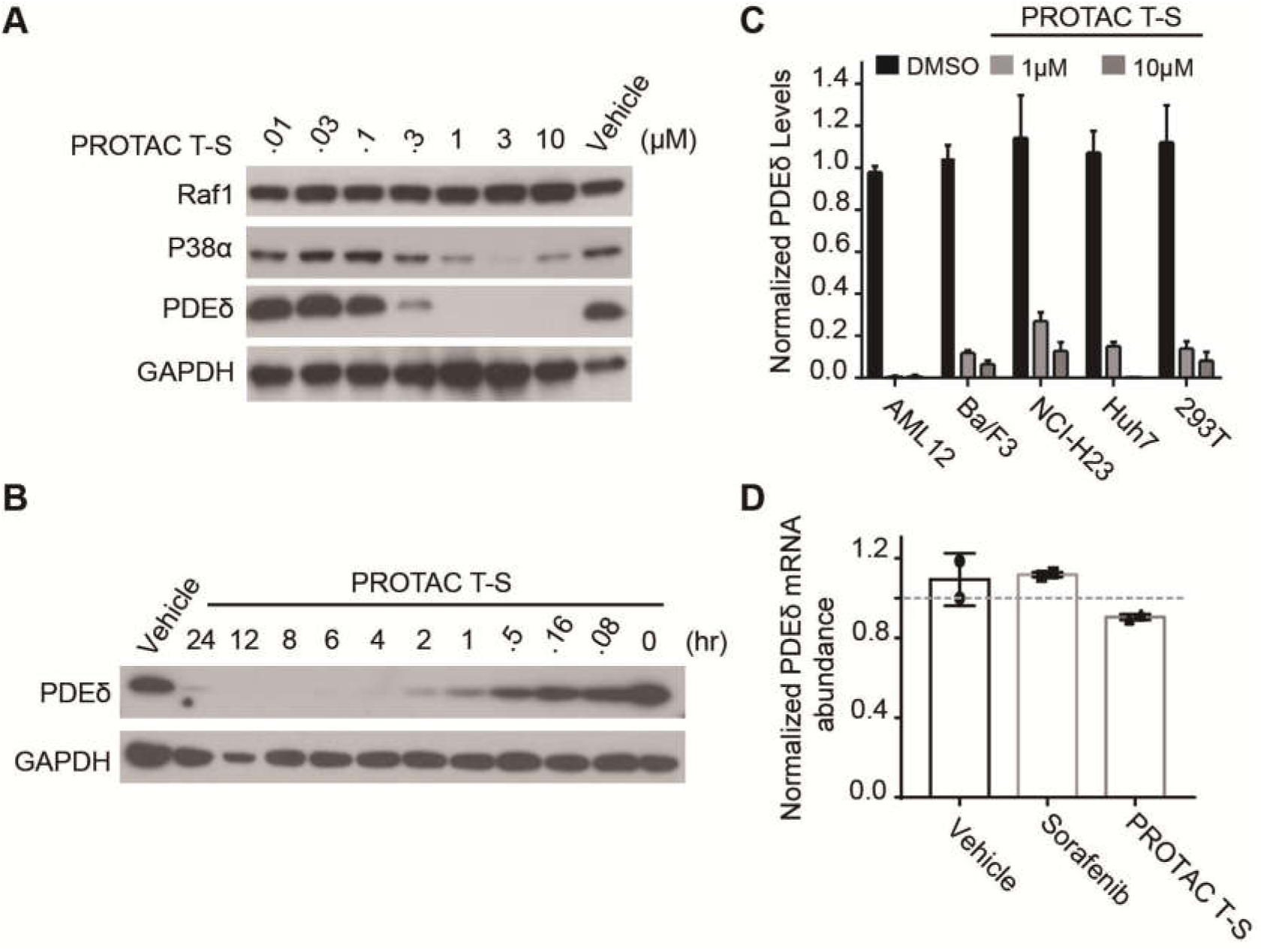
PDEδ was potently degraded by PROTAC T-S. (A) Western blot analysis of Raf1, P38α and PDEδ protein levels in AML12 in the presence of PROTAC T-S. (B) Western blot analysis of time-dependent degradation of PDEδ protein in the presence of 2 μM PROTAC T-S in 293T cells. (C) Degradation of PDEδ mediated by 1 μM or 10 μM PROTAC T-S in different cell lines. (D) Quantitative real-time PCR was performed after 12 h of treatment with either vehicle (DMSO), sorafenib (2 μM) or PROTAC T-S (2 μM) in AML12 cells. Data were normalized with GAPDH. Error bars denote s.d (n=2).

### 2.3. PROTAC T-S-induced degradation of PDEδ depends on ternary complex formation and ubiquitin-proteasome system

To determine whether PROTAC T-S induced PDEδ degradation through the formation of ternary PDEδ:PROTACT-S:CRBN complex, we used thalidomide and sorafenib to compete with PROTAC T-S to dissociate the ternary complex. Both thalidomide and sorafenib treatments could reduce the degradation of PDEδ mediated by PROTAC T-S (Figure 3A). The blockage of PDEδ degradation was more significant after a combined treatment of thalidomide and sorafenib, but the amount of PDEδ protein (about 52%) still could not return to untreated base levels even at high concentration of thalidomide and sorafenib. Using a high affinity PDEδ binding compound deltarasin, we saw the complete blockage of PROTAC T-S induced PDEδ degradation. Deltarasin treatment alone has no effect on PDEδ expression levels. Similar results were also observed in CRBN knock-out cells, where PDEδ protein was stable even at high concentration of PROTAC T-S (Figure 3B). To further confirm the formation of ternary PDEδ:PROTACT-S:CRBN complex, we tested whether PROTAC T-S could induce ubiquitination of PDEδ in the cellular system. When 293T cells co-expressing FLAG-PDEδ and HA-Ub were treated with vehicle or 500 nM PROTAC T-S for 1 hour, only the PROTAC T-S-treated cells displayed substantial ubiquitination of immunoprecipitated FLAG-PDEδ (Figure 3C). It was expected that the inhibition of ubiquitin-proteasome system could block the degradation of PDEδ induced by PROTAC T-S. Western blotting analysis showed that the degradation of PDEδ was totally blocked after pretreatment with proteasome inhibitor (MG132, bortezomib or carfilzomib) or neddylation inhibitor (MLN4924) (Figure 3D). These results strongly demonstrated that PROTAC T-S-mediated degradation of PDEδ fully depends on the formation of a ternary PDEδ:PROTAC T-S:CRBN complex and through the ubiquitin-proteasome system.

**Figure 3.**
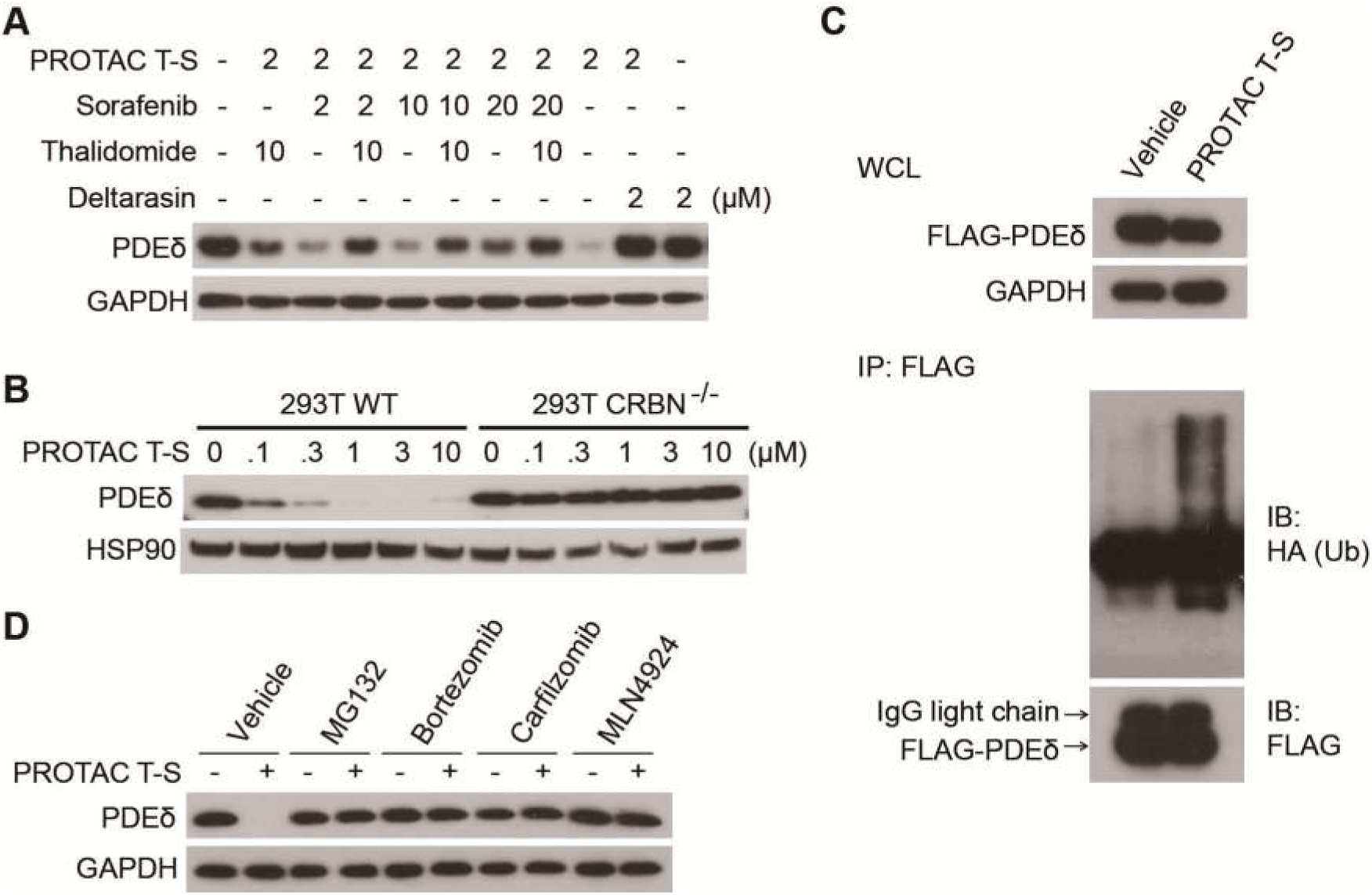
PROTAC T-S degradation of PDEδ is dependent on ternary complex formation and ubiquitin-proteasome system. (A) Western blot analysis of PDEδ protein in 293T cells. 293T cells were treated with PROTAC T-S for 8 hours in the presence or absence of the indicated dose of sorafenib, thalidomide or deltarasin. (B) Western blot analysis of PDEδ protein in 293T wildtype and CRBN knockout cells in the presence of PROTAC T-S. (C) Western blot analysis of PDEδ protein in AML12 cells. AML12 cells were treated with DMSO, MG-132 (10 μM), bortezomib (0.4 μM), carfilzomib (0.4 μM) or MLN4924 (1 μM) for 1 h and subsequently treated with either vehicle (DMSO) OR PROTAC T-S (2 μM) for 5 h. (D) PROTAC T-S ubiquitinates FLAG-PDEδ in 293T cells. 293T cells co-transfected with HA-Ubiquitin and FLAG-PDEδ were subsequently treated with either vehicle (DMSO) or PROTAC T-S (2 μM) for 1 h. Ubiqutinated FLAG-PDEδ was assessed via western blots detecting HA-Ubiquitin.

Of note, the degradation efficacy of PROTAC T-S toward P38α was easily and largely abolished by the competition of thalidomide or sorafenib (Figure S4), suggesting that the PDEδ:PROTAC T-S:CRBN complex is more stable than the P38α:PROTAC T-S:CRBN complex. Considering that the PROTAC T-S induced a more efficient degradation of PDEδ than P38α (Figure 2), our results are consistent with previous studies showing that the more potent the ternary complex formation, the more efficient the target degradation^5, 16, 29^.

### 2.4. PDEδ is a non-kinase target of sorafenib

To further confirm that PDEδ is also a direct target of sorafenib in cells, we used the CETSA assay to test whether PDEδ could be stabilized by sorafenib. Deltarasin as the high affinity PDEδ binding ligand strongly increased the stability of PDEδ in 293T cells even at 67°C (Figure 4). The stability of PDEδ was also significantly increased in the presence of sorafenib. Sorafenib treatment induced obvious shifts in the melting curve of PDEδ compared to the vehicle group. Thus, the CETSA experiments indicated that PDEδ is the target that is engaged by sorafenib in cells. Furthermore, we analyzed the interaction between sorafenib and PDEδ by using surface plasmon resonance (SPR) (Table 1). Sorafenib and PROTAC T-S showed similar binding affinity to PDEδ (1.23 μM and 2.26 μM, respectively). Therefore, we concluded that PDEδ is a novel non-kinase sorafenib target.

**Table 1.**
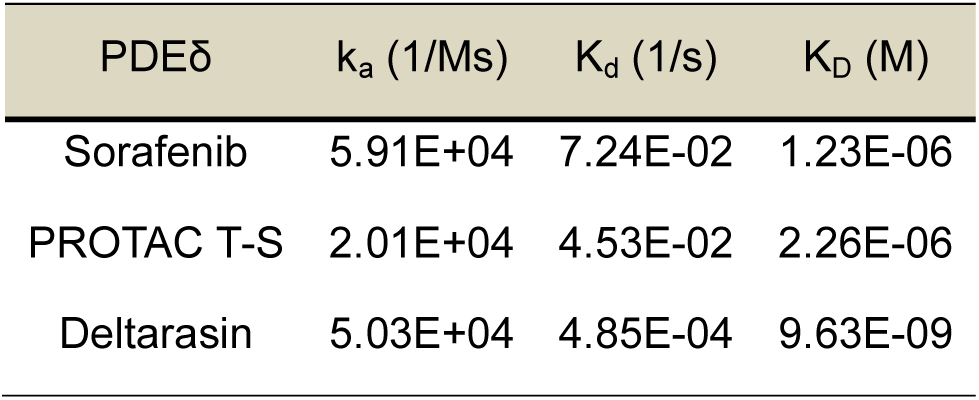
SPR results showed sorafenib, PROTAC T-S and deltarasin bind with PDEδ.

| PDEδ | $k_a$ (1/Ms) | $K_d$ (1/s) | $K_D$ (M) |
| --- | --- | --- | --- |
| Sorafenib | 5.91E+04 | 7.24E-02 | 1.23E-06 |
| PROTAC T-S | 2.01E+04 | 4.53E-02 | 2.26E-06 |
| Deltarasin | 5.03E+04 | 4.85E-04 | 9.63E-09 |

**Figure 4.**
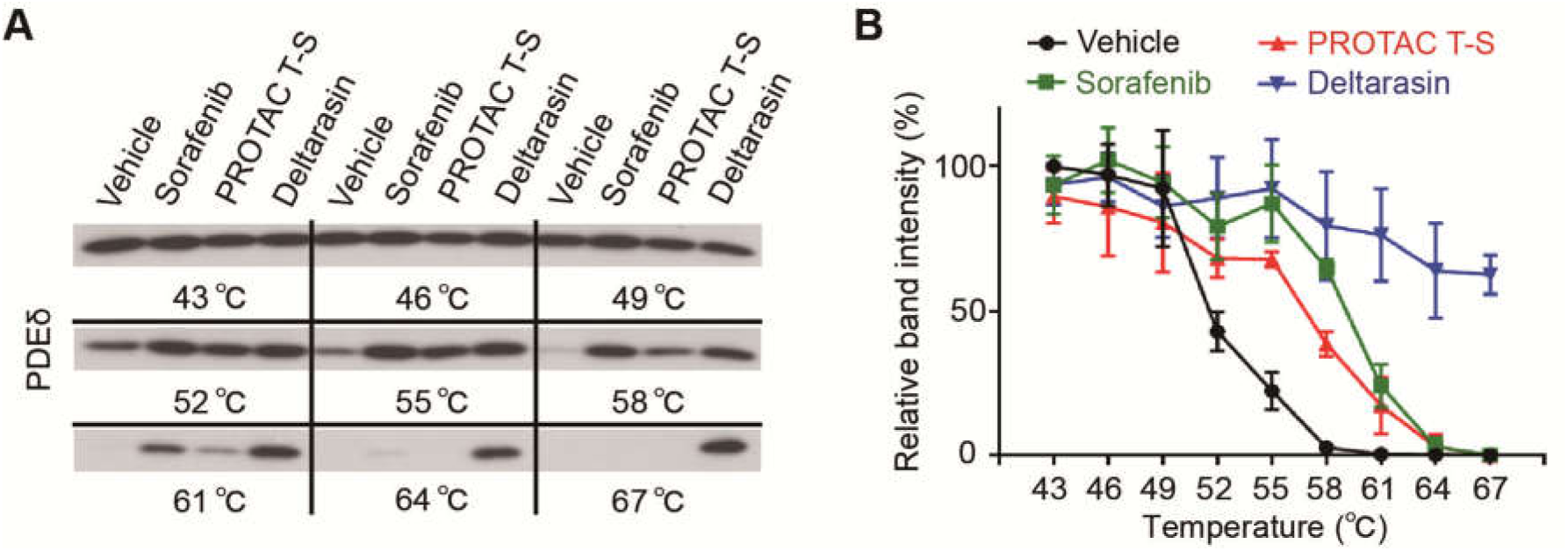
PDEδ protein levels were stabilized by sorafenib. (A) Western blot analysis of PDEδ protein levels in the presence of sorafenib, PROTAC T-S or deltarasin (positive control) treatment. 293T cells treated with 20 μM compounds for 2 hours and subsequently heated at different temperatures (43-67°C). (B) Quantification of PDEδ protein levels based on western blots from A. Error bars denote s.d (n=3).

Recently, PDEδ was reported to be overexpressed in sorafenib-resistant HCC cells, and contributed to migration and proliferation of liver cancer^30^. Therefore, PROTAC T-S could be used in overcoming the sorafenib resistance in such cancer cells. Although it is beyond the scope of our current study, it is desirable to further elucidate the role of PDEδ in sorafenib-produced anticancer effect and antifibrotic activity, as such information may facilitate the clinical usage of sorafenib as well as designing new drugs with dual inhibition activities of kinases and PDEδ.

### 2.5. Sorafenib binding mode in PDEδ revealed by molecular docking

To further validate the binding interactions between sorafenib and PDEδ, we docked sorafenib into the PDEδ binding pocket. According to the model, sorafenib is deeply buried into the hydrophobic pocket of PDEδ (Figure 5A). The urea oxygen and methylformamide oxygen atoms of sorafenib formed hydrogen bonds with Arg61 and Tyr149 of PDEδ, respectively. Moreover, Arg61, Tyr149 and the benzimidazole nitrogens of deltarasin formed two hydrogen bonds, which were critical for PDEδ-deltarasin binding, had been found in PDEδ-deltarasin crystal structure^24^. To validate our proposed model, we mutated Tyr149 to alanine (Y149A) to disrupt the hydrogen bond. As expected, PDEδ Y149A mutation protein was no longer degraded in the presence of PROTAC T-S (Figure 5B). Since the methylpicolinamide nitrogen atoms critical for the kinase binding did not form any specific interactions in our docking model, we used an in-house tool compound to further validate our model, HN02, previously designed to remove the kinase activities of sorafenib (Figure S5)^3^. The stability of PDEδ, but not P38α, showed an obvious increase in the presence of HN02 compared with sorafenib by CETSA (Figure 5C-D and Figure S6). These results showed that oxygen atom but not methylpicolinamide nitrogen of sorafenib takes part in the interaction with PDEδ.

**Figure 5.**
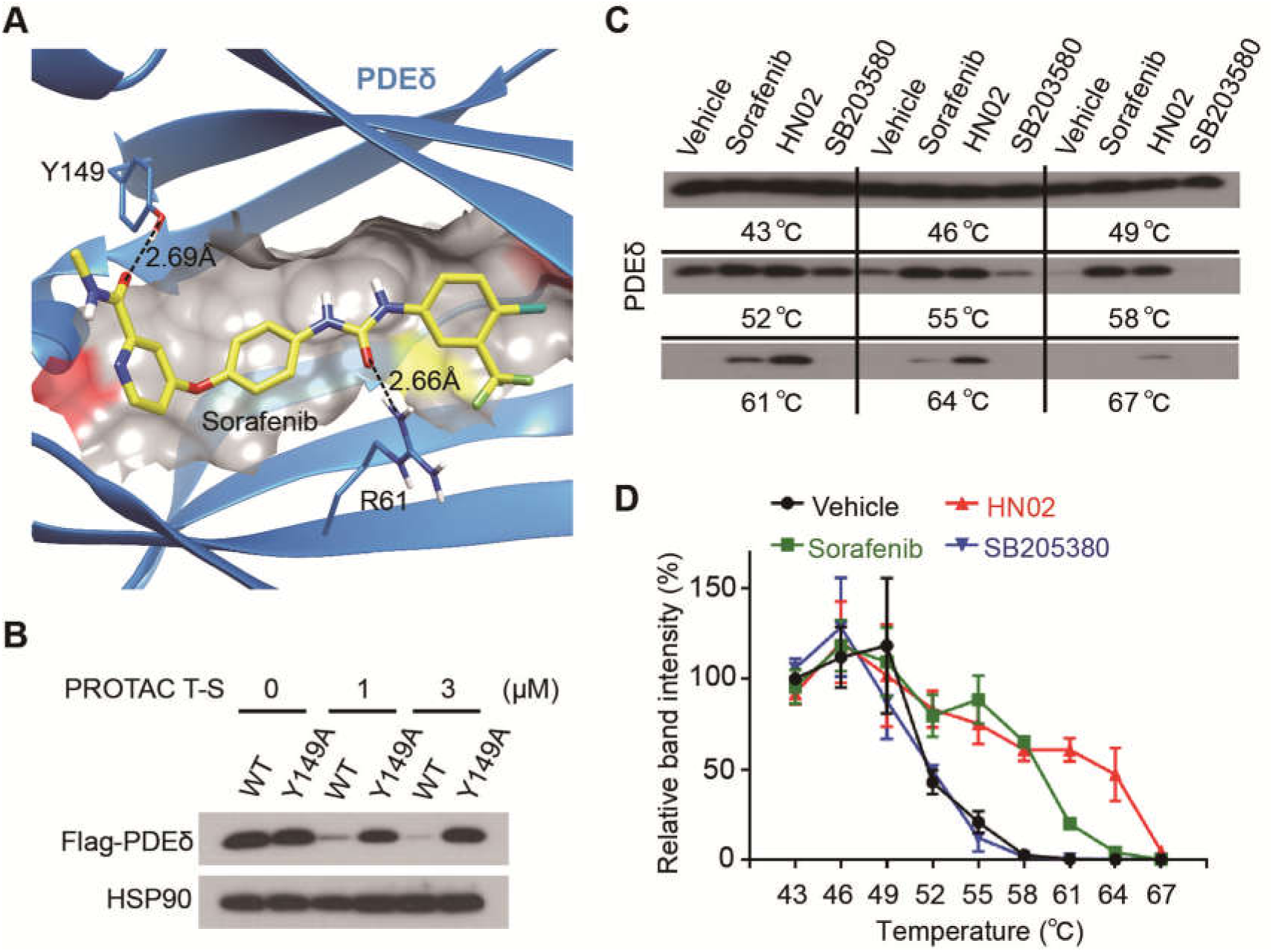
The binding mode of the PDEδ-sorafenib complex. (A) Molecular docking structures of sorafenib (yellow) onto PDEδ (blue, PDB code 3T5I). Arginine 61 and tyrosine 149 on PEδ formed two hydrogen bonds with sorafenib. Molecular images were generated with UCSF Chimera. (B) Western blot analysis of overexpressed PDEδ WT or Y149A mutant protein levels in 293T cells in the presence of PROTAC T-S. HSP90 was used as the loading control. (C) Western blot analysis of PDEδ protein level in the presence of sorafenib, HN02 or SB203580 treatment. 293T cell treated with 20 μM compounds for 2 hours and subsequently heated at different temperatures (43-67°C). (D) Quantification of PDEδ protein levels based on western blot from C. Error bars denote s.d (n=3).

### 2.6. PROTAC technology complements traditional approach in drug target identification

In addition, we used the traditional affinity-based proteomics approaches by biotinylation of sorafenib to purify the potential sorafenib targets in cells (Figure S7A). After excluding targets that had also been identified in control groups, a total of 61 proteins were found that uniquely interact with biotin-sorafenib (Figure S7B). We only found ATXN10, but not PDEδ or other previously identified sorafenib’s targets, in the list. To confirm that ATXN10 is also a target of PROTAC T-S, we tested the ATXN10 protein levels by Western blotting and found it was largely reduced in the presence of PROTAC T-S (Figure S7C). Therefore, ATXN10 could be another potential target of sorafenib. The reason for the low overlap of the identified targets between PROTAC-based and affinity-based approaches, probably due to the biotin modification, affected the binding affinity of sorafenib, but this was compensated in PROTAC by formation of the ternary complex. Although we did not further validate the other three degraded proteins (BCCIP, MAP3K20, and MAPK2) found by MS screening, it is very likely that two protein kinases MAP3K20 and MAPK2 are potential target of sorafenib. Therefore, we believe that PROTAC-based approach can be used as an alternative method for drug target identification, which is shown to have superior signal-to-noise ratio, in striking contrast to other affinity or activity-based approaches. However, arguably, PROTAC-based approach does suffer from other drawbacks such as missing potential targets due to relying on the engagement of ternary Target:PROTAC:E3 ligase complex.

## 3. Conclusions

In this study, we report the design and synthesis of sorafenib-based multi-target degrader PROTAC T-S. By combining this with quantitative proteomics, we identified non-kinase targets PDEδ was degraded by PROTAC T-S and could be potential targets of PROTAC T-S and sorafenib. The degradation of PDEδ mediated by PROTAC T-S was via the ubiquitin-proteasome system. PROTAC T-S could be a powerful compound to study the function of PDEδ in cells. The direct binding interaction between PDEδ and sorafenib was confirmed using CETSA and SPR assays. Furthermore, we dissected the molecular basis of sorafenib-PDEδ interaction, which happens through the methylformamide oxygen atom (but not nitrogen atom which was seen in sorafenib-kinase crystal structures), forming a hydrogen bond with the Tyr149 of PDE6δ. Altogether, our results identify a new target of sorafenib and potentially provide a novel perspective for further understanding of the mechanism of action of sorafenib. In addition, PROTAC-based approach expands the chemical proteomics toolbox to identify unknown targets of clinical drugs.

## 4. Experimental

### 4.1. Chemistry

All reagents and organic solvents were commercially available and used without further purification. ^1^H NMR, ^19^F NMR were measured on a Varian Inova-400 Spectrometer with tetramethylsilane (TMS) as internal standard. Data for ^1^H NMR spectra are reported relative to CDCl_3_ (7.26 ppm), or DMSO-*d*_6_ (2.50 ppm) as an internal standard and are reported as follows: chemical shift (δ ppm), multiplicity (s = singlet, d = doublet, t = triplet), coupling constant *J* (Hz), and integration. Mass spectra data were obtained on an ACQUITY UPLC system. All tested compounds had a purity >95%.

#### 1.1.1. 4-hydroxythalidomide (2)

Compound **2** was prepared from 4-nitroisobenzofuran-1,3-dione, and the Sandmeyer reaction was according to the established paper^31^. The subsequent process was modified. Before the sublimation, anhydrous Na_2_SO_4_ (equal mass) was added, then it was sublimated under reduced pressure with strong stirring. Contrasted to the formal sublimation, this improvement enhanced the yield from 51.3% to 86.3% for two steps. Then it was reacted with 3-amino-2,6-dioxo-piperidine hydrochloride in a sealed tube at 120°C, which afforded 3-hydroxythalidomide as a white solid in 78.2% yield. ^1^H NMR (400 MHz, DMSO-*d*_6_) δ 11.17 (s, 1H), 11.09 (s, 1H), 7.65 (dd, *J* = 8.4, 7.2 Hz, 1H), 7.34 – 7.29 (m, 1H), 7.25 (d, *J* = 8.2 Hz, 1H), 5.07 (dd, *J* = 12.8, 5.4 Hz, 1H), 2.88 (ddd, *J* = 17.3, 14.0, 5.4 Hz, 1H), 2.64 – 2.51 (m, 2H), 2.07 – 1.96 (m, 1H).

#### 1.1.2. tert-butyl 2-((2-(2,6-dioxopiperidin-3-yl)-1,3-dioxoisoindolin-4-yl)oxy)acetate (3)

A flame-dried 50 mL flask with Teflon stir bar, was charged with **2** (1.37 g, 5.0 mmol, 1.0 eq) and K_2_CO_3_ (1.04 g, 7.5 mmol, 1.5 eq) in DMF (20 mL), then tert-Butyl bromoacetate (970 mg, 5.0 mmol, 1.0 eq) was dropwised at room temperature. The mixture was stirred at room temperature for 3 h. After completion, the reaction mixture was diluted with EA (40 mL), washed with saturated NaCl solution (40 mL×3). The organic phase was separated, dried over Na_2_SO_4_, concentrated to afford crude product. The crude product was purified by silica chromatography with EA/PE(1/3), which afforded product as a white solid (1.36 g, yield: 70%). ^1^H NMR (400 MHz, DMSO-*d*_6_) δ 11.11 (s, 1H), 7.80 (dd, *J* = 8.5, 7.3 Hz, 1H), 7.48 (d, *J* = 7.2 Hz, 1H), 7.38 (d, *J* = 8.5 Hz, 1H), 5.10 (dd, *J* = 12.8, 5.4 Hz, 1H), 4.97 (s, 2H), 2.95 – 2.87 (m, 1H), 2.64 – 2.54 (m, 2H), 2.06– 2.02 (m, 1H), 1.43 (s, 9H).

#### 1.1.3. 2-((2-(2,6-dioxopiperidin-3-yl)-1,3-dioxoisoindolin-4-yl)oxy)acetic acid (4)

A flame-dried 50 mL flask with Teflon stir bar, was charged with **3** (390 mg, 1.0 mmol, 1.0 eq) in TFA/DCM (5 mL/20 mL). The mixture was stirred at room temperature for 2 h. The solvent was removed under reduced pressure, which afforded product as a white solid (332 mg, quantitive). ^1^H NMR (400 MHz, DMSO-*d*_6_) δ 11.08 (s, 1H), 7.76 (dd, *J* = 8.5, 7.3 Hz, 1H), 7.44 (d, *J* = 7.2 Hz, 1H), 7.36 (d, *J* = 8.5 Hz, 1H), 5.07 (dd, *J* = 12.8, 5.4 Hz, 1H), 4.96 (s, 2H), 2.90 – 2.82 (m, 1H), 2.53 (dd, *J* = 21.4, 11.0 Hz, 2H), 2.05 – 1.95 (m, 1H)

#### 1.1.4. tert-butyl(1-((2-(2,6-dioxopiperidin-3-yl)-1,3-dioxoisoindolin-4-yl)oxy)-2-oxo-7,10,13-trioxa-3-azahexadecan-16-yl)carbamate (5)

A flame-dried 50 mL flask with Teflon stir bar, was charged with **4** (167 mg, 0.5 mmol, 1.0 eq) and *tert*-butyl (3-(2-(2-(3-aminopropoxy)ethoxy)ethoxy)propyl)carbamate (320 mg, 1.0 mmol, 2.0 eq) in EtOH (8 mL), then DMT-MM (276 mg, 1.0 mmol, 2.0 eq) was added in portions at room temperature. The suspension was stirred at room temperature for 1 h. After completion, the solvent was removed under reduced pressure, and the residue was purified by flash system with MeCN/H_2_O, which afforded product as a light yellow oil (288 mg, yield: 91% including *tert*-butyl (3-(2-(2-(3-aminopropoxy)ethoxy)ethoxy)propyl)carbamate). ^1^H NMR (400 MHz, CDCl_3_) δ 7.74 (dd, *J* = 8.3, 7.4 Hz, 1H), 7.61 – 7.52 (m, 2H), 7.19 (d, *J* = 8.3 Hz, 1H), 4.97 (dd, *J* = 12.2, 5.5 Hz, 2H), 4.68 – 4.57 (m, 2H), 4.07 – 4.01 (m, 4H), 3.68 – 3.39 (m, 15H), 3.19 (d, *J* = 6.1 Hz, 2H), 2.82 (dddd, *J* = 13.4, 11.6, 9.2, 4.7 Hz, 3H), 2.15 (ddd, *J* = 7.7, 6.3, 3.6 Hz, 1H), 1.91 – 1.83 (m, 2H), 1.74 (dd, *J* = 12.3, 6.1 Hz, 6H), 1.42 (d, *J* = 4.0 Hz, 9H).

#### 1.1.5. N-(3-(2-(2-(3-aminopropoxy)ethoxy)ethoxy)propyl)-2-((2-2,6-dioxopiperidin-3-yl)-1,3-dioxoisoindolin-4-yl)oxy)acetamide (6)

A flame-dried 50 mL flask with Teflon stir bar, was charged with **5** (64 mg, 0.1 mmol, 1.0 eq) in TFA/DCM (1 mL/4 mL). The mixture was stirred at room temperature for 2 h. The solvent was removed under reduced pressure, which afforded product as a white solid (54 mg). It was used directly for the next step without further purification.

#### 1.1.6. 4-(4-(3-(4-chloro-3-(trifluoromethyl)phenyl)ureido)phenoxy)-N-(1-((2-(2,6-dioxopiperidin-3-yl)-1,3-dioxoisoindolin-4-l)oxy)-2-oxo-7,10,13-trioxa-3-azahexadecan-16-yl)picolinamide (PROTAC T-S)

A flame-dried 15 mL flask with Teflon stir bar, was charged with **6** (16 mg, 0.03 mmol, 1.0 eq) and 4-(4-(3-(4-chloro-3-(trifluoromethyll)phe-nyl)ureido)phenoxy)picolinic acid (13 mg, 0.03 mmol, 1.0 eq) in EtOH (4 mL), then DMT-MM (17 mg, 0.06 mmol, 2.0 eq) was added in portions at room temperature. The suspension was stirred at room temperature for 1 h. After completion, the solid was filtered out, and the filtrate was concentrated under reduced pressure. The residue was purified by flash system with MeCN/H_2_O, which afforded product as a white solid (18 mg, yield: 61% for 2 steps). ^1^H NMR (400 MHz, CD_3_OD) δ 8.43 (d, *J* = 5.6 Hz, 1H), 7.97 (d, *J* = 2.6 Hz, 1H), 7.76 (dd, *J* = 8.4, 7.4 Hz, 1H), 7.62 (dd, *J* = 8.6, 2.6 Hz, 1H), 7.56 – 7.51 (m, 2H), 7.49 (dd, *J* = 9.0, 1.6 Hz, 3H), 7.38 (d, *J* = 8.0 Hz, 1H), 7.12 – 7.06 (m, 2H), 7.02 (dd, *J* = 5.6, 2.6 Hz, 1H), 5.10 (dd, *J* = 12.6, 5.5 Hz, 1H), 4.71 (s, 2H), 3.64 – 3.50 (m, 10H), 3.48 – 3.42 (m, 2H), 3.38 (t, *J* = 6.7 Hz, 2H), 3.34 – 3.31 (m, 2H), 3.24 (dd, *J* = 3.3, 1.6 Hz, 1H), 2.82 (dd, *J* = 13.3, 4.4 Hz, 2H), 2.72 (dd, *J* = 18.6, 6.1 Hz, 3H), 2.12 (s, 1H), 1.80 (ddd, *J* = 19.2, 12.7, 6.5 Hz, 4H). Purities of PROTAC T-S determined by LC-MS were >95% (shown in Supporting Information).

### 4.2. Kinase Inhibition Assay

The enzymatic activities of BRAF, c-KIT, FLT3, VEGFR2 and PDGFRβ were performed at Reaction Biology Corporation following standard protocols. Sorafenib and PROTAC T-S were treated in single dose duplicate mode at a concentration of 0.1 µM. The enzymatic activity of p38α MAPK was performed as previously described^32^. Nonspecific inhibition was prevented by adding 0.1% (v/v) Triton X-100 in P38α kinase activity assay.

### 4.3. Cell Culture and Stable Cell Line Generation

AML12 cell line (Typical Culture Preservation Commission Cell Bank, Shanghai) was maintained in DME/F-12 1:1 (Hyclone) supplemented with ITS Liquid Media Supplement (Sigma). HEK293T (ATCC) and Huh7 (ATCC) cell lines were maintained in DMEM (Gibico). NCI-H23 (ATCC) cell line and Ba/F3 cells stable expressing K-Ras G12D (kindly provided by Dr. Liang Chen, Jinan University) were maintained in RPMI 1640. All cells were cultured at 37 °C with 5% CO_2_. For CRBN knock out cell lines, on day one, AML12 or 293T cells were seeded in 6 well plates, and co-transfected the cells next day with Cas9 plasmide, gRNA plasmids (5′-CGCCCGTCCCGGAGGTTACC-3′ for mouse and 5′-AACTACTCCGGGCGGTTACC-3′ for human) and EGFP-expressing plasmid at a ratio of 3:2:1 when the cells were grown to 85-90% confluence. Two days later, the GFP-positive cells were sorted into single cell by flow cytometry (BD Biosciences FACSAria II). The individual clones were verified by DNA sequencing or Western blotting.

### 4.4. Western Blot

Cells were seeded in 6-well or 12-well plates, after treated by indicated dose of compounds the cells were washed twice by cold phosphate-buffered saline (PBS) and lysed in RIPA lysis buffer (Sigma) containing protease inhibitor (Roche). Equal amount proteins of each sample were separated by SDS-PAGE and transferred to polyvinylidene difluoride (PVDF) membrane. The PVDF membranes were first blocked in TBS-T (Tris-buffered saline containing 0.1% Tween 20, pH 7.5) containing 5% skim milk and then incubated with specific antibodies. Finally, the blot signal was detected by ECL substrate (Millipore) and captured by using X-Ray film (Nikon).

### 4.5. Mass Spectrometry for Proteomics

AML12 cells were treated by PROTAC T-S or Thalidomide for 12 hours, then lysed with lysis buffer (150 mM NaCl, 1% NP-40, 0.25% sodium deoxycholate and 1% sodium dodecyl sulfate, pH 7.4) containing 1mM PMSF and protease inhibitor (Roche). The lysates were sonicated (5 × 6 s) and centrifuged at 20,000 × g for 20 min at 4 °C. The supernatant fraction of the cell extract was collected, and the protein concentration was quantified by BCA assay (Thermo Fisher Scientific). The proteins were reduced using 10 mM tris(2-carboxyethyl)phosphine (TCEP) for 30 minutes at room temperature, then alkylated in the dark for 30 minutes using 50 mM iodoacetamide. Above proteins was precipitated with pre-chilled acetone and centrifuged at 20,000 × g for 10 minutes at 4°C. Precipitated protein pellets was resuspended with 100 mM triethylammonium bicarbonate (TEAB) buffer (pH 8.5) and digested by trypsin (1: 50) overnight.

Tandem Mass Tag (TMT) 10-plex isobaric labeling were performed as manufacturer’s instructions. After labeling, the peptides from the six samples were pooled together in equal proportion. The pooled TMT samples were fractionated using off-line, high-pH reverse-phase (RP) chromatography on an Xbridge BEH C18 column (4.6 × 250 mm, 3.5um, Waters) on a Agilent 1290 system. The samples were separated using a 40-min gradient of solvents A (10 mM formate in water, pH 9) and B (10 mM ammonium formate in 80% acetonitrile, pH 9), at a flow rate of 1 mL/min. Peptides were separated into 40 fractions and then consolidated into 10 fractions.

All fractions were analyzed sequentially on Orbitrao Fusion (Thermo Fisher Scientific) equipped with Dionex RSLCnano HPLC (Thermo Fisher Scientific). Peptides were injected onto a EasySpray column (75 μm × 25 cm, PepMap RSLC C18 column, 2 μm, 100 Å, Thermo Fisher Scientific). A mix of buffer A (0.1% formic acid in water) and B (0.1% formic acid in 80% acetonitrile) was used over a linear gradient from 8 to 35% buffer B over 90 min using a flow rate of 300 nL/min. The spray was initiated by applying 2.3 kV to the EASY-Spray emitter and the data were acquired under the control of Xcalibur software in a data-dependent mode using top speed and 3s duration per cycle. The survey scan is acquired in the orbitrap covering a range from 375-1500 m/z at resolution of 120,000 and an automatic gain control (AGC) target of 4.0 x 105 ions, Maxium Injection Time 50 ms. The most intense ions were selected for fragmentation using HCD in Orbitao with 38% collision energy and an isolation window of 1.4 Th. The fragments were then analysed in the orbitrap with a resolution of 60,000. The AGC target was set to 1.0 x 105 and the maximum injection time was set to 105 ms with a dynamic exclusion of 60 s, The raw data files for all 10 fractions were merged and searched against the mouse protein database (uniprot-taxonomy-10090) by Maxquant for protein identification and TMT reporter ion quantitation. The identifications were based on the following criteria: enzyme used trypsin; maximum number of missed cleavages equal to two; precursor mass tolerance equal to 10 ppm; fragment mass tolerance equal to 0.02 Da; variable modifications: oxidation (M); fixed modifications: carbamidomethyl (C). The data was filtered by applying a 1% false discovery rate followed by exclusion of proteins with less than two unique peptides. Quantified proteins were filtered if the absolute fold-change difference ≥1.42.

### 4.6. Protein Expression and Purification

The human PDEδ cDNA contained a stop code was cloned into the pET-28a vector. PDEδ with an N-terminal 6× His was produced in E. coli BL21 (DE3) cells. Cells were grown at 37°C in LB medium containing 50 µg/mL of Kanamycin. His-tagged PDEδ protein was induced at 18°C by 0.5 mM IPTG overnight. Cell pellet was resuspended and soniced in Tris buffer (20 mM Tris-HCl, 150 mM NaCl, pH 8.0) containing 1 mM PMSF. His-tagged PDEδ protein in supernatant was purified using HisTrap column (GE Healthcare), HiTrap Q column (GE Healthcare) and Superdex 75 column (GE Healthcare).

### 4.7. Ubiquitination assays

The ubiquitination assay was performed as previously described^29^. Briefly, 293T cells were seeded in 6 cm dishes and allowed to adhere overnight. The nest day, cells were transfected with FLAG-PDEδ and HA-Ubiquitin by using Lipofectamine 2000 (Invitrogen). After 24 h, cells were treated with either vehicle (DMSO) or PROTAC T-S for 1 hour. Then cells were washed twice with ice-cold PBS and lysed in 500 µL cell lysis buffer (25 mM Tris-HCl, 150 mM NaCl, 1% NP-40, 1% sodium deoxycholate, 0.1% SDS, pH 7.6) containing protease inhibitor (Roche). The immunoprecipitation was performed according to the protocol of anti-DDDDK-tag mAb-magnetic agarose (MBL). Agarose were resuspended in lithium dodecyl sulfate (LDS) sample buffer containing 5% 2-mercaptoethanol (BME) and heated at 70 °C for 10 min. Finally, the ubiquitinated FLAG-PDEδ (anti-HA-tag) was evaluated by western blot.

### 4.8. Cellular Thermal Shift Assay

293T cells were treated with the test compounds (20 µM) or DMSO control for 2 hours in 10 cm dish, then washed twice with PBS, harvested and resuspended in PBS containing protease inhibitor (Roche), distributed 100 µL of cell suspensions into each 0.2 mL PCR tubes and heated at indicated temperatures (43-67°C) for 3 min. Finally, the tubes were incubated at room temperature for 3 min, followed by two freeze-thaw cycles to extract the total proteins. The soluble fraction was then analyzed by western blotting.

### 4.9. Surface Plasmon Resonance (SPR)

The binding affinity of PROTAC T-S, Sorafenib and deltarasin against PDEδ protein was measured via SPR technology using Biacore 8k instrument (GE Healthcare) at room temperature. The PDEδ protein in 10 mM sodium acetate (pH 5.5) was covalently immobilized on biosensor chips (CM5) by amine coupling. The experiment was performed in PBS running buffer supplemented with 0.05% Tween 20 and 5% DMSO. The compounds were injected in five concentrations (2-fold serial dilution) using the injection at 30 µL/min flow rate. In the dissociation phase, the buffer flowed for 180s. The binding kinetics and binding affinity of those compounds were analyzed using Biacore 8K Evaluation Software (GE Healthcare).

### 4.10. Prediction binding pose of sorafenib/PDEδ

Molecular docking is applied to predict the binding pose of sorafenib and PDEδ. Four structures of PDEδ binding to different ligands were selected as receptors (PDB: 3T5I, 4JVF, 5E80 and 5ML4). Then sorafenib was docked into receptor binding pockets by DOCK3.7^33^. For each docking simulation, up to 20 binding poses were saved except for receptor 5E80, in which only one binding pose was generated due to limited pocket volume. In total 71 predicted poses were rescored following the previously described.^34^ The best rescored pose is used for following study.

### 4.11. Biotinylated-sorafenib Pull-down Assay

Firstly, streptavidin-sepharose beads (GE Healthcare) were washed three times with PBS and incubated with 500 μM biotin or biotinylated-sorafenib for 8 h at 4 °C. Then the beads were washed three times with wash buffer (50 mM Tris-HCl, 150 mM NaCl, 1% Triton X-100, pH 7.4) and incubated with the supernatants of AML12 cells lysis. One tube of biotinylated-sorafenib containing beads was added with high dose sorafenib to compete the specific sorafenib-binding proteins. After incubated overnight with gentle rotation at 4 °C, the beads were washed three times with wash buffer and boiled for 5 min in 40 μL SDS loading buffer. The supernatant was separated by a SDS-PAGE and stained with silver stain reagents (Sigma). The total gel strips were excised for mass spectrometry analysis.

## Acknowledgments

We thank Dr. She Chen in the proteomics center at the National Institute of Biological Sciences, Beijing for assistance with LC−MS/MS experiments. Financial supports from the Chinese Ministry of Science and Technology “973” Grant 2014CB849802 (to N.H.),Central Public-interest Scientific Institution Basal Research Fund 2017ywf-zd-16 (to Q.M.) and National Natural Science Foundation of China Grant 21807007 (to Y.Z.) are gratefully acknowledged. Computational support was provided by Special Program for Applied Research on Super Computation of the NSFC-Guangdong Joint Fund (the second phase) under Grant U1501501.

## Author contributions

Yang Li, Qingshi Meng, Xiangbing Qi and Niu Huang conceived the project. Yang Li, Qingshi Meng, Ping Wang, Xiaolei Liu, Qiuyu Fu, Yuting Xie and Yu Zhou performed the experiments. Yang Li and Niu Huang analyzed the data and wrote the manuscript.

## Conflicts of interest

The authors have no conflicts of interest to declare.

## Appendix A.

Supporting Information

---

**Scheme 1.**
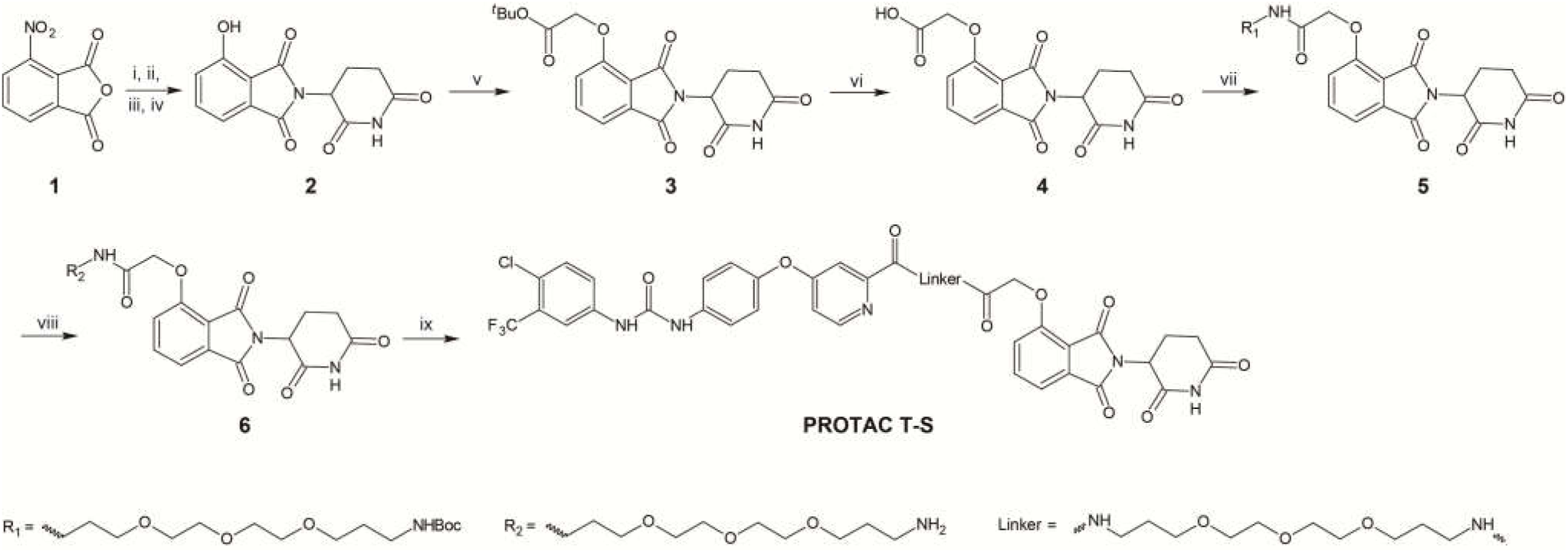
General procedure for PROTAC T-S synthesis^*a*^. ^*a*^Conditions: (i) 10% Pd/C (dry), molecular sieve (4A), H_2_ (balloon), EA, room temperature, 3 h, 89%; (ii) H_2_SO_4_, NaNO_2_, room temperature, 4h; (iii) 0.02 KPa, Na_2_SO_4_, sublimation, 117-141 °C, 86% (for two steps); (iv) 3-Amino-2,6-dioxo-piperidine hydrochloride, pyridine, 120 °C, 14 h, 78%; (v) *tert*-Butyl bromoacetate, K_2_CO_3_, DMF, room temperature, 3 h, 70%; (vi) TFA/DCM, room temperature, 2 h, quantitative; (vii) DMT-MM, EtOH, room temperature, 1 h, 91%; (viii) TFA/DCM, room temperature, 2 h; (ix) DMT-MM, EtOH, room temperature, 1 h, 61% for 2 steps.

## Notes

### Competing Interest Statement

The authors have declared no competing interest.

